# Early cadmium responses in developing oat caryopses indicate an unexpected regulatory network linked to low grain cadmium accumulation

**DOI:** 10.64898/2026.08.17.745199

**Authors:** Bitz Lidija, Bitz Oliver, Haikka Hanna, Hautsalo Juho, Tenhola-Roininen Teija, Tanhuanpää Pirjo, Frank Panitz

## Abstract

Heavy-metal accumulation in cereal grains is becoming critical for European food safety, regulation and plant breeding. In the EU, Cd maximum levels in certain foodstuffs have been revised, including lowering or establishing limits for relevant food categories, while new maximum levels for nickel (Ni) have recently been introduced for several foodstuffs, including cereal categories, with limits for oats and selected cereals applying from 2026. Together, these developments create an urgent need to identify genetic and physiological mechanisms that reduce Cd and Ni accumulation in cereal grains while maintaining crop quality and productivity. Against this regulatory and food-safety background, our broader RNA-seq experiment investigates early transcriptional responses to Cd and Ni in oat F2 segregants contrasting for metal accumulation. The full dataset includes low- and high-accumulating segregants, roots and developing caryopses sampled at 3 h and 7 h after treatment. In the present pilot analysis, we focus on the Cd response in developing caryopses of the low-Cd accumulating segregant AS131 to identify candidate processes associated with reduced grain Cd accumulation. The strongest transcriptional responses were not dominated by canonical Cd-detoxification pathways. At 3 h after Cd exposure, differentially expressed transcripts were mainly associated with cell-wall functions, endosperm transfer-cell-specific PR60 proteins, DUF239-containing proteins and cysteine proteinase inhibitors, whereas several dehydration-, pathogen-, defence-, cell-wall-loosening- and ROS- related genes were repressed. By 7 h, the response suggested a shift towards homeostatic acclimation, with induction of TIP2 aquaporins, thiamine thiazole synthases, EF-Tu proteins, coatomer-related genes and carbohydrate metabolism-associated genes, together with repression of LEA/SMP/dehydrin genes, FRO7-like genes, EF-hand calcium-binding proteins and stress-regulatory transcription factors. Pathway-level analyses were broadly consistent with these transcript-level patterns, highlighting structural, nucleosome-associated, translation-related, metabolic and developmental processes. Several Cd-responsive transcripts were also associated with broader abiotic-stress responses, suggesting recruitment of shared stress-regulatory modules rather than Cd-specific detoxification pathways alone. Overall, these results support a working hypothesis in which low Cd accumulation in developing oat grain may involve regulation of solute-transfer interfaces, cellular protection, intracellular homeostasis, trafficking pathways and caryopsis developmental programmes. These findings provide candidate processes for future comparison with high-Cd accumulating segregants, root tissues and Ni responses in the broader dataset.

## Introduction

Cadmium (Cd) is a non-essential toxic metal that enters the human diet primarily through food, with cereals representing a major exposure source because of their high consumption (Apuli et al., 2025). Reducing grain Cd accumulation remains important for food safety and regulatory compliance and will require complementary agronomic and genetic solutions (Apuli et al., 2025; Mei et al., 2022). This challenge is particularly relevant for oat (Avena sativa L.), an increasingly important component of Nordic diets (Blomhoff et al., 2023), where grain Cd concentrations may exceed accepted limits under some conditions.

Grain Cd accumulation is a complex trait governed by metal uptake, translocation, redistribution, sequestration and detoxification. These processes involve transporter families such as ZIP, NRAMP, CDF, YSL, ABC and P1B-type ATPases, together with metal-chelating compounds including phytochelatins, metallothioneins and nicotianamine (Colangelo and Guerinot, 2006; Cobbett and Goldsbrough, 2002; Haydon and Cobbett, 2007; Mei et al., 2022; Tsednee, 2024). Major advances have been achieved in several cereals. In durum wheat, the Cd locus Cdu1 was traced to TdHMA3-B1, where a non-functional transporter allele increases grain Cd concentration (Maccaferri et al., 2019). In rice, transporters such as OsNRAMP5, OsHMA2 and OsHMA3 play key roles in Cd uptake, translocation and sequestration, while in maize ZmHMA3 has been implicated in grain Cd accumulation (Mei et al., 2022; Tang et al., 2021). These studies demonstrate the importance of transporter-mediated regulation, while also highlighting species-specific genetic architectures (Tsednee, 2024).

In oat, a major QTL affecting grain Cd concentration was previously mapped in an F₂ population derived from the cultivars Aslak and Salo (Tanhuanpää et al., 2007). Although this established a strong genetic basis for variation in grain Cd accumulation, the underlying gene(s) and the transcriptional responses associated with Cd accumulation remain unresolved. Transcriptomic studies in ryegrass, maize and wheat have successfully identified Cd-responsive genes, pathways and regulatory candidates (Hu et al., 2020; Liu et al., 2021; Li et al., 2022), but direct comparison among studies is complicated by differences in species, tissues, developmental stages and experimental designs.

The broader RNA-seq experiment included low- and high-accumulating F₂ segregants exposed to Cd and Ni, with roots and developing caryopses sampled at 3 h and 7 h after treatment. Here, we focus on the early Cd response in developing caryopses of the low-Cd accumulating segregant AS131. We aimed to identify transcriptional responses and candidate biological processes associated with Cd exposure and to determine whether the response was dominated by canonical metal-detoxification pathways or by broader transport, homeostatic, metabolic and developmental processes. By integrating GO enrichment, homology-based pathway analyses and transcript-level investigation, this study provides a hypothesis-generating framework for understanding early Cd responses in developing oat grain.

## Materials and methods

### Plant material

In the study of Eurola et al. (2003) it was found out that Swedish spring hulled oat cultivar «Salo» (Svalöf-Weibull AB) was high in accumulating Cd from the fields in Finland (mean 0.060 mg kg-1 of dry weight being more than a twice much than the lowest observed). For developing molecular markers linked to the Cd accumulation a F2 population containing 150 individuals has been developed from a cross Aslak × Salo, Aslak being Finnish spring hulled cultivar (Boreal Plant Breeding Ltd.). Aslak was tested to be low Cd accumulator (roots, leaves and grains) when compared to Salo with the most significant difference coming from the grains (Tanhuanpää et al., 2007). The distribution of the Cd concentration in grains of the Aslak × Salo F2 progeny fit the hypothesis of single gene inheritance with the dominant allele for low accumulation having some of the phenotypes more extreme than the parents suggesting that some minor genes also influenced Cd accumulations. Tanhuanapää et al. (2007) mapped QTL affecting Cd accumulations representing a major gene for low grain Cd concentration.

To advance gene studies on low cadmium (Cd) accumulation, we selected the F2 genotype AS131 from Tanhuanpää et al. (2007) for this pilot study on differentially expressed genes (DEGs). This genotype was identified as one of the nine that exhibited the lowest Cd accumulation (>5x fold after 0.5mg/kg Cd treatment i.e. lower than the low parent Aslak, 650 µg seed versus 3490 µg). The AS131 seeds for this experiment were obtained from Natural Resources Institute Finland (Luke) (Jokioinen, Finland) where they are stored and maintained.

### Greenhouse experimental design, cadmium treatment, time points and sampling

One seed of the low-Cd accumulating Aslak × Salo F2 segregant AS131 was sown per 3.5 L pot containing 1050 g of peat-soil mixture. A total of fifteen AS131 plants were grown in a greenhouse at 17 °C/15 °C day/night temperature under a 16 h photoperiod.

Cadmium treatment was selected in relation to reported Cd concentrations in Finnish soils, ranging from 0.02 to 0.75 mg kg^−1^ (Mäkelä-Kurtto et al., 2003). At 14 days post anthesis (DPA), Cd was applied to the pot substrate at 1 mg kg^−1^ as Cd(NO_3_)_2_ × 4H_2_O (Fluka) dissolved in water. Each plant received 200 mL of Cd solution, applied both from above and to the saucer below the pot. Regular watering was omitted on the treatment day to promote absorption of the Cd solution, which was taken up within approximately 10 min.

Because AS131 consisted of F2 progeny and individual plants were not genetically fixed, pre-treatment caryopses were collected from the same plants immediately before Cd application and used as controls. This within-plant control design was chosen to minimise genetic variation between control and Cd-treated samples. After Cd application, developing caryopses were collected from the same plants at 3 h and 7 h after treatment and stored in liquid nitrogen.

Three biological replicates were collected for each sampling point: pre-treatment control, 3 h after Cd treatment and 7 h after Cd treatment. Each replicate was obtained from a different plant grown in a separate pot. Approximately two developing caryopses from the upper part of the panicle were pooled per plant to obtain one RNA sample. In total, nine RNA-seq samples were prepared per plant. The parental cultivars Aslak and Salo were grown in parallel under the same experimental conditions to enable comparison of Cd concentration patterns between the parental genotypes and AS131. In addition, untreated control plants were included to establish baseline Cd concentrations.

### Cadmium concentration in mature grains

Mature seeds from the panicles of all plants were harvested and replicates from the same time points were merged into one sample to determine\validate the Cd levels before and after Cd treatment. At the same time parents were grown and cadmium measured. The seeds were dried at 65 °C, pulverized and Cd content was measured by inductively coupled plasma mass spectrometry (ICP-MS) (PerkinElmer ELAN 6000) as described by (Eurola et al., 2003).

### RNA isolation and quality assessment

Collected caryopses were immediately frozen in liquid nitrogen and stored at −80 °C until RNA extraction. Total RNA was extracted using the PowerPlant RNA Isolation Kit (Qiagen, Germany), followed by RNase-free DNase treatment (Qiagen) to remove residual genomic DNA. RNA was eluted in 100 µL of buffer and assessed for integrity, purity and concentration.

RNA integrity was evaluated using an Agilent 2100 Bioanalyzer with RNA chips. RNA purity was assessed using a NanoDrop One/OneC spectrophotometer (Thermo Fisher Scientific) based on absorbance ratios at 260/280 nm, and RNA concentration was determined using a Qubit 4 Fluorometer (Thermo Fisher Scientific). RNA samples with sufficient quantity (>1000 ng), high purity and high integrity (RIN > 8.5) were used for cDNA library preparation and Illumina RNA sequencing.

### Library construction and RNA-seq

For each RNA-seq sample, 100 ng of high-quality total RNA was used for library preparation. Libraries were prepared from three biological replicates per treatment and time point using the QuantSeq 3′ mRNA-Seq Library Prep Kit FWD for Illumina (Lexogen, Vienna, Austria), according to the manufacturer’s instructions with individual indexes per sample Equal amounts of RNA were used from pre-treatment control samples and from the corresponding Cd-treated AS131 samples collected at 3 h and 7 h after treatment.

Libraries were pooled and sequenced at the Natural Resources Institute Finland (Luke, Jokioinen, Finland) using an Illumina NextSeq 550 system with a NextSeq 500/550 High Output v2.5 kit. Sequencing was performed as single-end 1 × 75 bp reads, targeting more than 20 million reads per sample for gene expression profiling. Raw reads were subjected to quality control before downstream expression analysis.

### Data analysis

Fastp (v0.23.4) (Chen et al., 2018) was used to remove potential Illumina Universal adapters (“AGATCGGAAGAG”) from raw sequences and to trim ‘length_required’ to 50 bp. Reads representing ribosomal RNAs were filtered using SortMeRNA (v4.3.6, smr_v4.3_default_db) (Kopylova et al., 2012). Quality assessments were visualised using FastQC and reports from read processing across the samples were generated with MulitQC (v.1.19).

As reference genome Avena sativa cv Sang v1.1 from EnsemblPlant (release 58, January 2024)(Yates et al., 2022) was used. We quantified the expression of transcripts with the quasi-mapper Salmon (v1.9.0; Patro et al., 2017); with parameters --seqBias –validateMappings -- libType=SF) in mapping-based mode using a decoy-aware transcriptome index. This index uses the transcriptome targets in the reference as well as the whole genome as decoys and was built with a k of 31 for our reads of about 74bp. The library type for QuantSeq FWD data is determined by the stranded single-end protocol where the reads come from the forward strand. The Salmon quantification result quant.sf files contain the information on read counts which will be used for differential expression (DE) analysis.

### Differential expression analysis

Post-processing analyses were performed using R Statistical Software (version 4.3.1) and the tidyverse package (Wickham et al., 2019). Exploratory data analysis (data not shown) was performed using the vsn package (version 3.68.0) (Huber et al., 2002) for variance stabilisation, pheatmap (version 1.0.12) (Kolde, Raivo, 2019) for heat maps of sample distances and clustering. For data visualisation the package ggplot2 (version 3.4.3) (Wickham, 2009) was used for plotting, while knitr (version 1.44) (Xie, 2015) created reports.

The Bioconductor package tximport (Soneson et al., 2015) is applied to import transcript-level abundance from Salmon quantification. Essentially, as the counts from 3’ tagged RNA-seq data do not have length bias no gene-level summarization is applied, instead tximport setting txOut=TRUE is used to output transcript-level data, and the original counts as counts matrices. The Bioconductor package DESeq2 package (version 1.40.2) (Love et al., 2014) was used to test for differential expression of genes. As design for the DESeqDataSetFromMatrix we used batch correction for the sequencing runs and a variable combining treatment (control, treatment) and time (0 h, 3 h, 7 h). The results from the DESeq2 DE analysis are obtained as contrasts between samples treated per time point vs control sample. Minimal pre-filtering was applied to ensure that at least 3 samples had a read count of at least 10 reads. For each time point treated samples (3 h and 7 h post-treatment) were compared against the control (pre-treatment). The detection and treatment of count outliers was performed by DESeq2 as previously described (Love et al., 2014). The automatic independent filtering provides multiple testing adjustment using the Benjamini–Hochberg method; the False Discovery Rate (FDR) cut-off alpha was set to 0.05 to identify differentially expressed genes.

### Annotation and enrichment analysis

For GO analysis/enrichment we used gProfiler (Kolberg et al., 2023) (accessed online 13 June 2024) with Avena sativa Sang as reference genome, using the differentially expressed transcripts identified in each sample. Pathway enrichment was performed with ShinyGO (Ge et al., 2020) (v0.80, accessed online 13 June 2024), however, since Avena sativa Sang genome was not available as reference we used orthologue genes from in dicot (Arabidobsis thaliana) and monocot (Oryza sativa Japonica Group) model species. Orthologues gene IDs for Avena sativa Sang transcript IDs were retrieved from EnsemblPlants BioMart (Kinsella et al., 2011; Yates et al., 2022). ShinyGO calculates enrichment P-values using hypergeometric test and corrects for multiple testing FDR is quantified using the Benjamini-Hochberg method; results are sorted by FDR, then by fold enrichment.

## Results

### Cadmium amount in the grains of the AS131 and parents Aslak × Salo

Seeds combined from different plants from mature Cd treated AS131, Aslak and Salo were picked, dried, grinded and Cd was measured using inductively coupled plasma mass spectrometry (ICP-MS) (Tab 1.).

### RNA sequencing

High-quality RNA from developing caryopses of the Aslak × Salo F2 low-Cd accumulating segregant AS131 was used for RNA-seq. Because AS131 consisted of non-fixed F2 progeny, pre-treatment caryopses from the same plants were used as controls to minimise genotype-related variation. After Cd application, caryopses were sampled from the same plants at 3 h and 7 h after treatment, with three biological replicates per sampling point.

Transcript abundance was quantified using Salmon. Reads assigned to annotated transcripts ranged from 52% to 58% across samples. This moderate assignment rate may partly reflect the use of a decoy-aware transcriptome index containing genome-wide decoy sequences, which reduces spurious assignment of reads from unannotated or genomic regions.

**Table 1.** Amount of Cd in the grains of Cd treated oats AS131, Aslak and Salo.

| Sample name | Cd Final result mg/kg | Cd concentration, subsample 1 (mg kg<sup>-1</sup>) | Dry matter, subsample 1 (%) |
| --- | --- | --- | --- |
| Aslak (0)* | 0.008 | 0.007 | 91.8 |
| Aslak | 0.088 | 0.080 | 91.4 |
| Salo | 0.436 | 0.398 | 91.2 |
| AS131 | 0.043 | 0.039 | 90.9 |

\*Aslak (0) indicates Aslak grown without Cd treatment.

### Global biological systems affected by Cd

Global GO analysis of all significant DEGs showed that structural molecule activity, protein heterodimerization, nucleosome-associated components and translation were the strongest overrepresented categories at both 3 h and 7 h (Tab. 2), whereas broad membrane-associated and kinase/signalling-related categories were underrepresented (Tab. 3). This suggests that the Cd-responsive transcriptome was dominated by cellular structural organization and protein-related processes rather than broad membrane-transport or signalling categories.

**Table 2.** Top five overrepresented GO terms among all significant DEGs at 3 h and 7 h after Cd treatment.

| Time point | Rank | GO ID | GO term | Interpretive note | Padj |
| --- | --- | --- | --- | --- | --- |
| 3h | 1 | GO:0005198 | Structural molecule activity | Structural/cellular organization; macromolecular structural components | $8.008 \times 10^{-136}$ |
| 3h | 2 | GO:0046982 | Protein heterodimerization activity | Protein-complex formation; protein interaction networks | $3.816 \times 10^{-91}$ |
| 3h | 3 | GO:0043232 | Intracellular non-membrane-bounded organelle | Non-membrane cellular structures; ribonucleoprotein/chromatin/cytoskeletal-associated organization | $2.599 \times 10^{-82}$ |
| 3h | 4 | GO:0000786 | Nucleosome | Chromatin organization; DNA packaging | $3.882 \times 10^{-74}$ |
| 3h | 5 | GO:0006412 | Translation | Protein synthesis; translational machinery | $1.046 \times 10^{-47}$ |
| 7h | 1 | GO:0005198 | Structural molecule activity | Structural/cellular organization; macromolecular structural components | $4.294 \times 10^{-65}$ |
| 7h | 2 | GO:0046982 | Protein heterodimerization activity | Protein-complex formation; protein interaction networks | $1.036 \times 10^{-48}$ |
| 7h | 3 | GO:0000786 | Nucleosome | Chromatin organization; DNA packaging | $1.507 \times 10^{-47}$ |
| 7h | 4 | GO:0043232 | Intracellular non-membrane-bounded organelle | Non-membrane cellular structures; ribonucleoprotein/chromatin/cytoskeletal-associated organization | $2.592 \times 10^{-37}$ |
| 7h | 5 | GO:0006412 | Translation | Protein synthesis; translational machinery | $2.819 \times 10^{-16}$ |

*GO enrichment was performed using all significant DEGs at each time point without separating upregulated and downregulated transcripts. Therefore, overrepresented terms indicate biological categories enriched among Cd-responsive genes.*

**Table 3.** Top five underrepresented GO terms among all significant DEGs at 3 h and 7 h after Cd treatment.

| Time point | Rank | GO ID | GO term | Interpretive note | Padj |
| --- | --- | --- | --- | --- | --- |
| 3h | 1 | GO:0016020 | Membrane | transmembrane transporter activity; signalling receptor activity | $8.363 \times 10^{-43}$ |
| 3h | 2 | GO:0043412 | Macromolecule modification | protein modification process; RNA modification; DNA modification | $8.098 \times 10^{-22}$ |
| 3h | 3 | GO:0004672 | Protein kinase activity | protein serine/threonine kinase activity; protein tyrosine kinase activity; protein histidine kinase activity | $1.489 \times 10^{-16}$ |
| 3h | 4 | GO:0032559 | Adenyl ribonucleotide binding | metabolic process; ATP binding; ABC transporter ATP-binding cassette domain binding | $3.472 \times 10^{-14}$ |
| 3h | 5 | GO:0043231 | Intracellular membrane-bounded organelle | nucleoplasm; nuclear chromosome; mitochondrial respiratory chain; mitochondrial matrix | $2.648 \times 10^{-12}$ |
| 7h | 1 | GO:0016020 | Membrane | transmembrane transporter activity; signalling receptor activity | $2.358 \times 10^{-14}$ |
| 7h | 2 | GO:0043412 | Macromolecule modification | protein modification process; RNA modification; DNA modification | $1.163 \times 10^{-11}$ |
| 7h | 3 | GO:0032559 | Adenyl ribonucleotide binding | metabolic process; ATP binding; ABC transporter ATP-binding cassette domain binding | $5.241 \times 10^{-9}$ |
| 7h | 4 | GO:0004672 | Protein kinase activity | protein serine/threonine kinase activity; protein tyrosine kinase activity; protein histidine kinase activity | $2.911 \times 10^{-7}$ |
| 7h | 5 | GO:0043231 | Intracellular membrane-bounded organelle | nucleoplasm; nuclear chromosome; mitochondrial respiratory chain; mitochondrial matrix | $5.606 \times 10^{-5}$ |

*GO enrichment was performed using all significant DEGs at each time point without separating upregulated and downregulated transcripts. Therefore, underrepresented terms indicate categories depleted among Cd-responsive genes. Underrepresentation should not be interpreted as transcriptional downregulation.*

#### Arabidopsis and rice homologue-based pathway enrichment of 7 h Cd-responsive oat genes

Homologue-based KEGG enrichment of 7 h Cd-responsive oat transcripts highlighted translation- and metabolism-associated pathways (Fig. 1). Arabidopsis homologues showed the strongest enrichment for ribosome-related pathways, together with carbohydrate, amino-acid, lipid/cuticle-related, ABC transporter, protein-processing and secondary-metabolism pathways (Fig. 1A). Rice homologues showed a similar pattern, with ribosome, arginine biosynthesis, amino-acid metabolism, photosynthesis, starch and sucrose metabolism, ribosome biogenesis and secondary metabolism among enriched pathways (Fig. 1B). These results provide supportive evidence that the 7 h response involved translation, metabolic adjustment and cellular maintenance.

**Figure 1.**
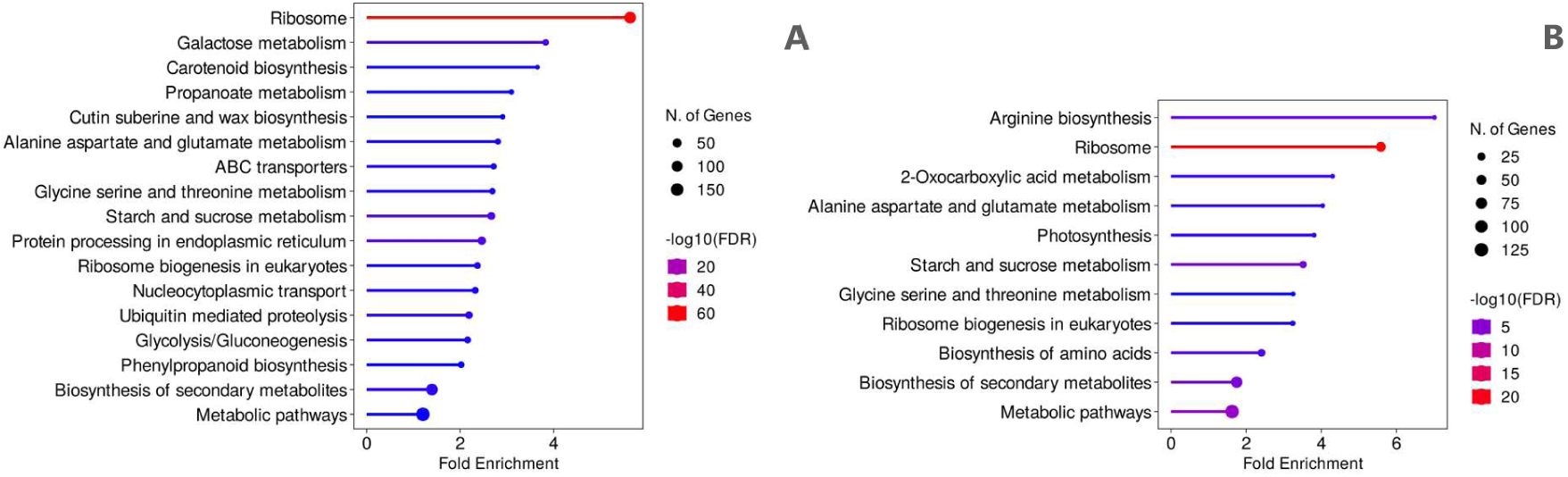
Homologue-based KEGG pathway enrichment of 7 h Cd-responsive oat genes using Arabidopsis and rice annotations. Oat DE transcripts detected 7 h after Cd treatment were mapped to homologous genes in A. thaliana and O. sativa japonica, and the corresponding gene identifiers were used for KEGG pathway enrichment. (A) Enriched KEGG pathways using A. thaliana homologues. (B) Enriched KEGG pathways using O. sativa japonica homologues. Fold enrichment is shown on the x-axis; point size indicates the number of genes assigned to each pathway; colour indicates −log10(FDR).

In addition, Plant Reactome and WikiPathways enrichment results were retained for the Arabidopsis homologue set as an additional supportive annotation layer (Fig. 2). These homologue-based analyses were interpreted cautiously and used only to provide broader functional context for the oat DEG results, recognizing that homology does not necessarily imply identical gene function.

**Figure 2.**
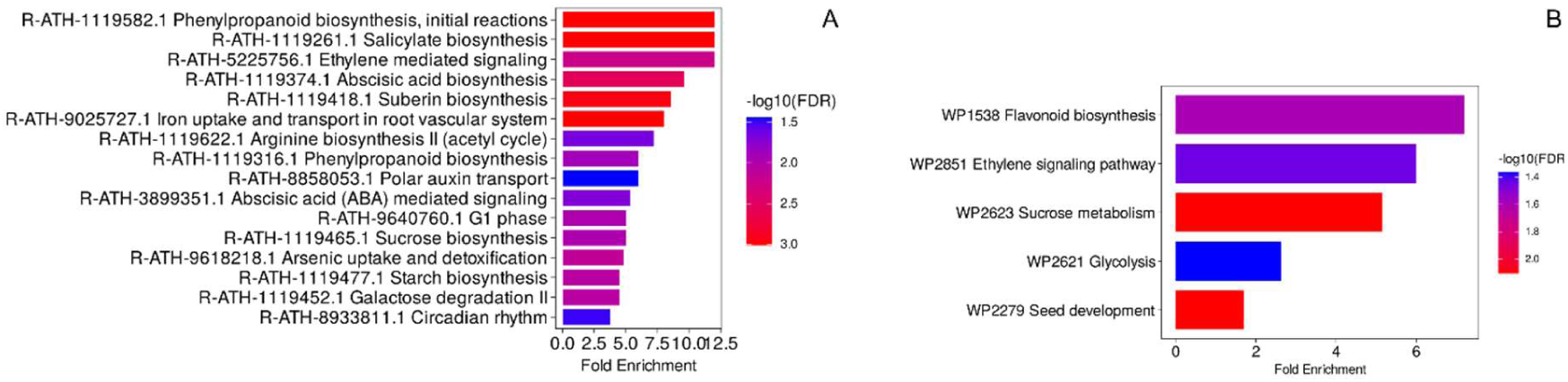
Homologue-based Plant Reactome and WikiPathways enrichment of 7 h Cd-responsive oat genes using Arabidopsis annotations. Oat DE transcripts detected 7 h after Cd treatment were mapped to homologous A. thaliana genes, and the corresponding Arabidopsis identifiers were used for pathway enrichment. (A) Enriched Plant Reactome pathways. (B) Enriched WikiPathways pathways. Fold enrichment is shown on the x-axis; colour indicates −log10(FDR).

### Differentially expressed genes underlying early response after Cd treatment

The differences in gene expression in AS131 developing caryopsis was investigated during three and seven hours after the Cd treatment compared to the control samples that were taken before Cd treatment at 0 h. Pre-treatment caryopses collected from the same AS131 plants immediately before Cd application were used as controls. A set of 1903 genes were differentially expressed 3 h after Cd treatment when compared to control samples (0h) (adj. p<0.05) out of which 1440 were upregulated and 463 downregulated (Tab. 4, Fig. 3A). At 7 h after Cd treatment, 1496 genes were differentially expressed (981 upregulated, 515 downregulated) (Tab 4., Fig. 3B). To enable detailed biological interpretation, we examined the most strongly responding transcripts within each response category.

**Figure 3:**
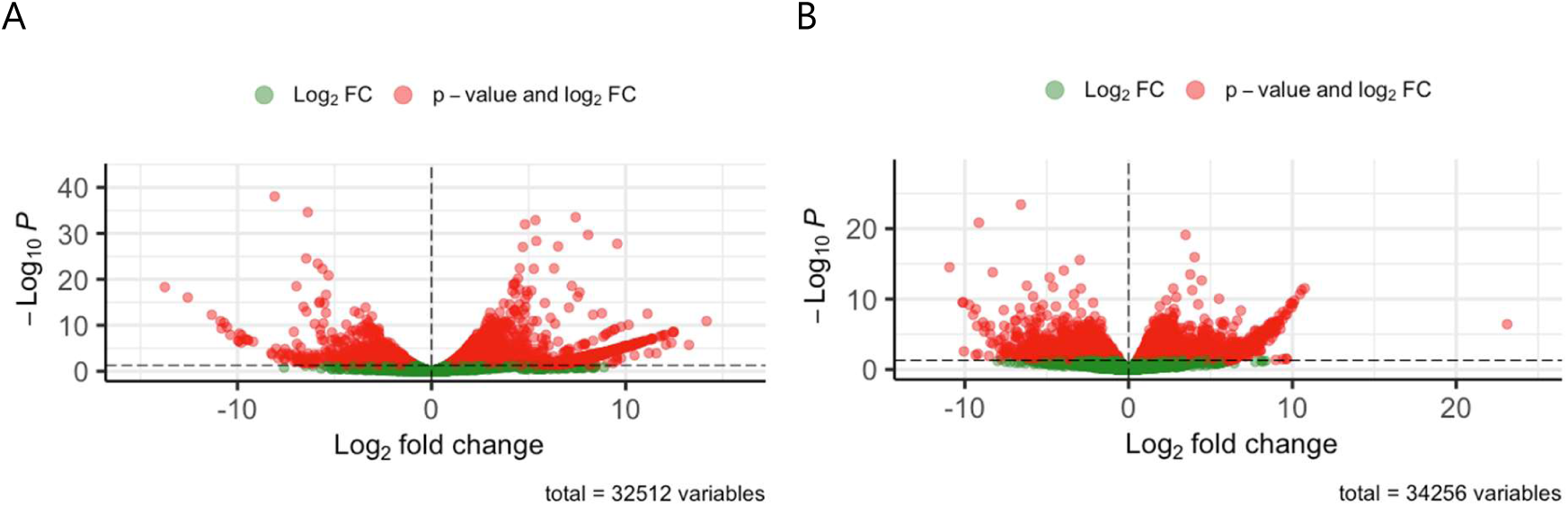
Differentially expressed genes in oat AS131 an Aslak × Salo F2 transgressive segreggant after 3 and 7 hours post Cd treatment when compared to control samples (0h before treatment). Volcano plots showing statistical significance vs magnitude of log2 fold change at 3 h (A) and at 7 h (B) after Cd treatment.

**Table 4:** The numbers of differentially expressed genes (DEGs) at 3 h and 7 h after Cd treatment.

|  | 3 h after Cd treatment<br>n(Padj)<0.05 | 7 h after Cd treatment<br>n(Padj)<0.05 |
| --- | --- | --- |
| Upregulated | 1440 | 981 |
| Downregulated | 463 | 515 |

n=number of genes

Candidate genes were not selected based on predefined functional categories. Instead, the most statistically significant upregulated and downregulated transcripts at each time point were inspected individually and subsequently organized into broader biological themes. To capture both statistically robust and strongly responsive transcripts, genes among the most significant (adjusted P-value) and those showing the largest expression changes (log2 fold change) were inspected. The top 10 differentially upregulated and downregulated genes at 3 h and 7 h after Cd treatment when considering both, adjusted p value and log fold change are shown in Tab. 5–8.

**Table 5.** Most strongly upregulated genes in developing caryopses of the low-Cd accumulating Aslak × Salo F2 segregant AS131 at 3 h after Cd treatment.

| Human readable description<br>Gene ID A. sativa Sang genome | Padj | Human readable description<br>Gene ID A. sativa Sang genome | Log2Fold<br>Change |
| --- | --- | --- | --- |
| cell wall protein precursor<br>AVESA.00010b.r2.2DG0352920.1 | 2.862E-30 | Glucose-1-phosphate adenylyltransferase<br>AVESA.00010b.r2.7CG0671800.1 | 14.17 |
| Endosperm transfer cell specific PR60<br>AVESA.00010b.r2.3AG0414350.1 | 9.757E-30 | Dihydrolipoyl dehydrogenase<br>AVESA.00010b.r2.3DG0560710.1 | 13.262 |
| Cysteine proteinase inhibitor<br>AVESA.00010b.r2.5AG0844240.1 | 6.409E-29 | Nucleosome assembly protein 1-like 1<br>AVESA.00010b.r2.1AG0039190.2 | 12.484 |
| NEP-interacting protein (DUF239)<br>AVESA.00010b.r2.5AG0837020.1 | 9.944E-27 | Pathogenesis-related thaumatin superfamily protein<br>AVESA.00010b.r2.3CG0510870.1 | 12.471 |
| Endosperm transfer cell specific PR60<br>AVESA.00010b.r2.3CG0461540.1 | 1.887E-25 | ZZ-type zinc finger-containing protein<br>AVESA.00010b.r2.2CG0305670.1 | 12.412 |
| cell wall protein precursor<br>AVESA.00010b.r2.2CG0272480.1 | 6.651E-25 | Endoglucanase<br>AVESA.00010b.r2.7AG1192400.1 | 12.356 |
| NEP-interacting protein (DUF239)<br>AVESA.00010b.r2.5AG0837040.1 | 2.181E-24 | Exportin-2<br>AVESA.00010b.r2.3DG0523410.1 | 12.083 |
| Endosperm transfer cell specific PR60<br>AVESA.00010b.r2.3AG0414380.1 | 2.558E-24 | Actin<br>AVESA.00010b.r2.1AG0061940.1 | 12.043 |
| Cysteine proteinase inhibitor<br>AVESA.00010b.r2.5CG0889690.1 | 8.554E-20 | Programmed cell death protein 4<br>AVESA.00010b.r2.2AG0227130.2 | 11.966 |
| Endosperm transfer cell specific PR60<br>AVESA.00010b.r2.3CG0461520.1 | 8.554E-20 | carboxyl-terminal peptidase (DUF239)<br>AVESA.00010b.r2.6AG1061820.1 |  |

**Table 6.**
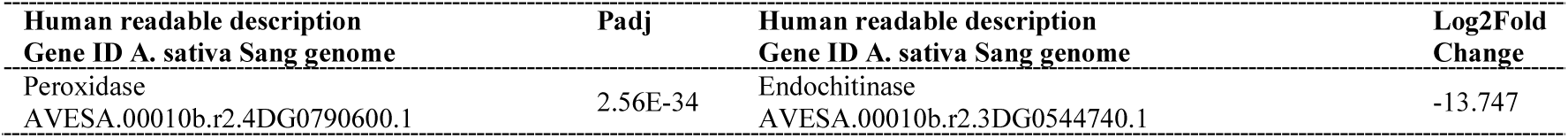
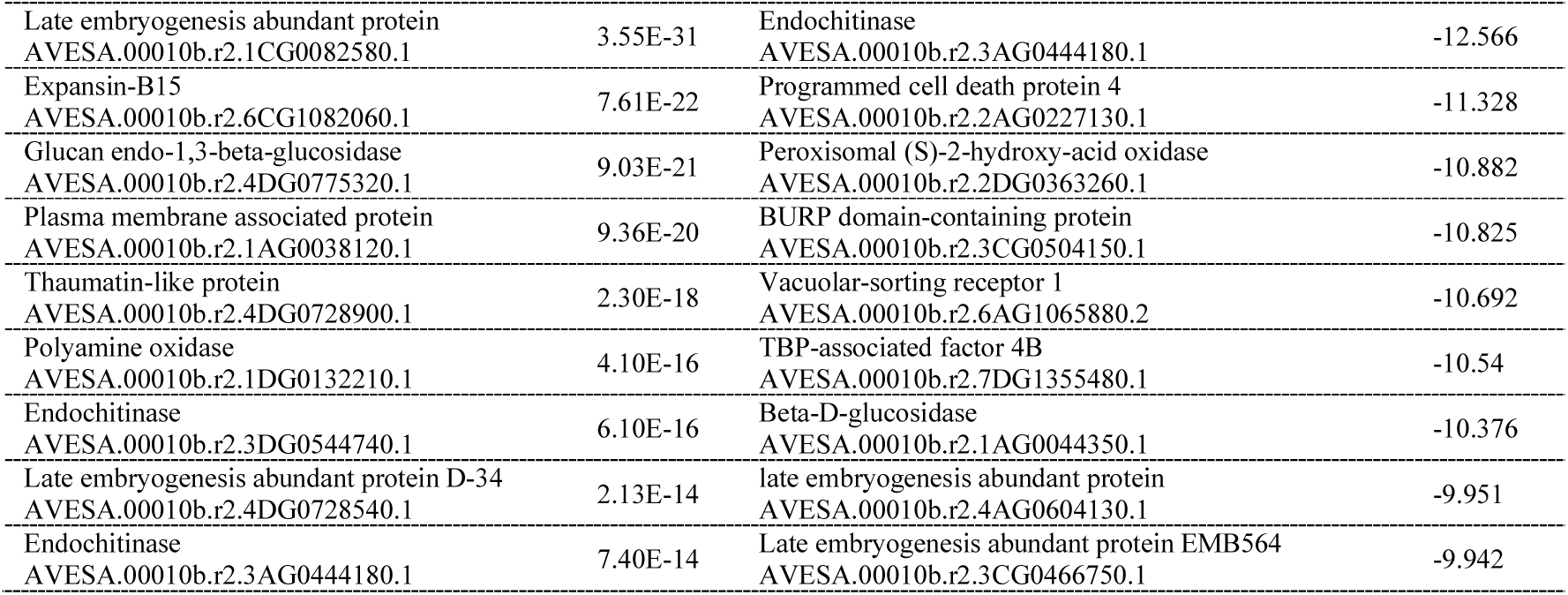
Most strongly downregulated genes in the low-Cd accumulating Aslak × Salo F2 segregant AS131 at 3 h after Cd treatment, ranked by log2 fold change and adjusted P value.

**Table 7.** Most strongly upregulated genes in the low-Cd accumulating Aslak × Salo F2 segregant AS131 at 7 h after Cd treatment, ranked by log2 fold change.

| Human readable description<br>Gene ID A. sativa Sang genome | Padj | Human readable description<br>Gene ID A. sativa Sang genome | Log2Fold<br>Change |
| --- | --- | --- | --- |
| tonoplast intrinsic protein 2<br>AVESA.00010b.r2.1AG0073470.1 | 8.595E-16 | Programmed cell death protein 4<br>AVESA.00010b.r2.2AG0227130.2 | 23.112 |
| tonoplast intrinsic protein 2<br>AVESA.00010b.r2.1DG0192290.1 | 9.258E-13 | GTP binding Elongation factor Tu family protein<br>AVESA.00010b.r2.4CG1257710.2 | 10.752 |
| Thiamine thiazole synthase, chloroplastic<br>AVESA.00010b.r2.2AG0197710.1 | 1.191E-10 | Hexosyltransferase<br>AVESA.00010b.r2.4DG0767710.3 | 10.563 |
| Thiamine thiazole synthase, chloroplastic<br>AVESA.00010b.r2.2DG0350970.1 | 6.8E-10 | Coatomer, beta' subunit<br>AVESA.00010b.r2.7DG1338400.2 | 10.465 |
| Cold induced 16<br>AVESA.00010b.r2.3DG0519330.1 | 6.612E-09 | Auxin efflux carrier family protein<br>AVESA.00010b.r2.5DG0958280.1 | 10.125 |
| GTP binding Elongation factor Tu family<br>protein<br>AVESA.00010b.r2.4CG1257710.2 | 6.612E-09 | casein kinase I<br>AVESA.00010b.r2.6CG1079270.1 | 10.051 |
| Hexosyltransferase<br>AVESA.00010b.r2.4DG0767710.3 | 1.038E-08 | Sucrose synthase<br>AVESA.00010b.r2.7DG1400080.3 | 9.954 |
| Histone H4<br>AVESA.00010b.r2.1DG0124280.1 | 1.038E-08 | Class I glutamine amidotransferase-like superfamily<br>protein<br>AVESA.00010b.r2.7AG1247170.1 | 9.95 |
| Coatomer, beta' subunit<br>AVESA.00010b.r2.7DG1338400.2 | 3.338E-08 | 60S ribosomal protein L9<br>AVESA.00010b.r2.5CG0867640.1 | 9.888 |
| tonoplast intrinsic protein 2<br>AVESA.00010b.r2.1CG0105710.1 | 8.024E-08 | Two-component response regulator ORR22<br>AVESA.00010b.r2.4DG0781390.1 | 9.727 |

only a selection among top 40 over-expressed genes were labeled on the volcano plots with gene ID

**Table 8.**
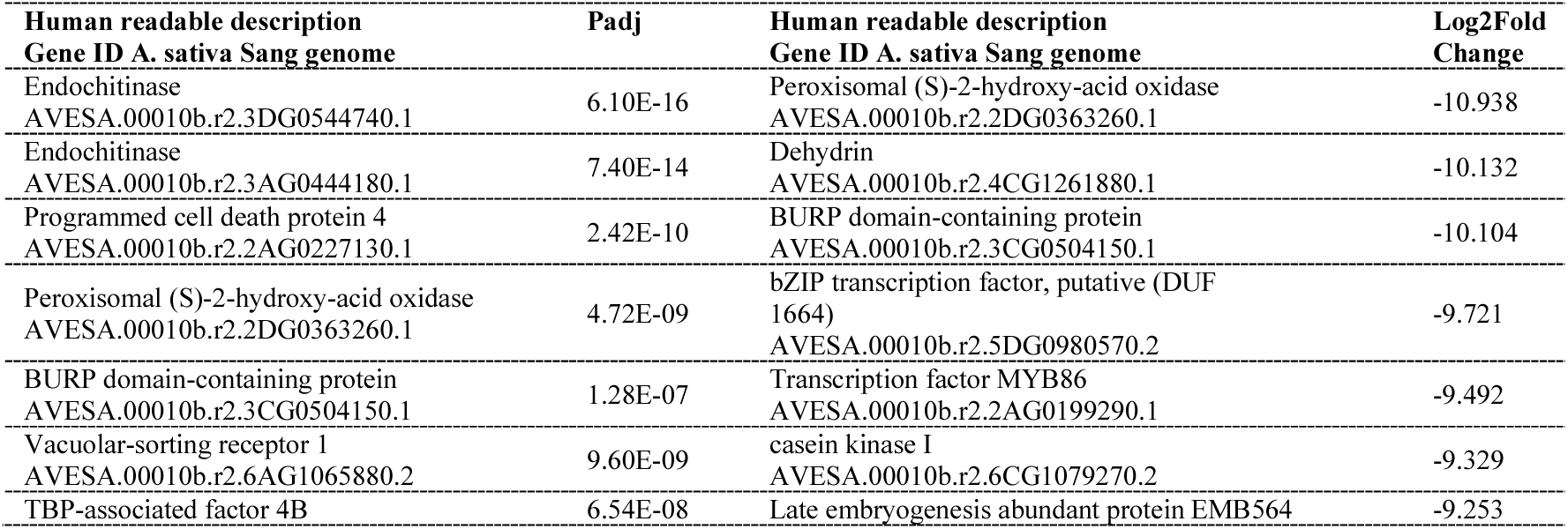
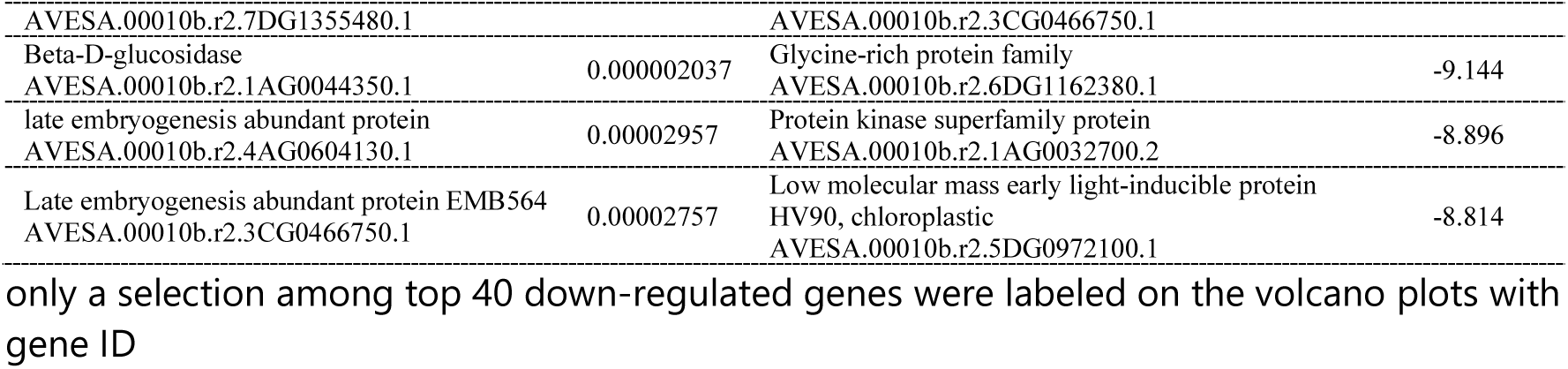
The most differentially down regulated genes according the log2fold change and adjusted p value in low accumulating F2 Aslak×Salo low segreggant AS131 at seven hours after Cd treatment.

## Discussion

### Cadmium accumulation in the grains of Aslak, Salo and and low Cd accumulating F2 progeny AS131

Mature grain Cd concentration in AS131 (0.043 mg kg⁻¹ DW) was lower than in Aslak (0.088 mg kg⁻¹ DW) and approximately tenfold lower than in Salo (0.436 mg kg⁻¹ DW), confirming the expected low-Cd phenotype (Tab. 1). Comparable values have been reported for other low-Cd cereal genotypes, such as wheat cv. Yaomai16 (0.066 mg kg⁻¹; Xiao et al., 2019). Although absolute Cd concentrations were lower than those reported by Tanhuanpää et al. (2007), likely due to the shorter Cd exposure period in the present study (14 dpa versus exposure from sowing), the relative ranking remained consistent across experiments, with AS131 showing the lowest and Salo the highest grain Cd accumulation.

### Enriched Biological Pathways

Functional enrichment was performed separately for all significant DEGs detected at 3 h and 7 h after Cd exposure, using unordered gene lists without filtering by log2 fold-change direction. Therefore, the analysis was used to identify biological categories broadly represented among Cd-responsive genes, rather than to infer pathway activation or repression.

At both time points, the most strongly overrepresented GO terms were associated with structural and protein-related cellular functions. Structural molecule activity, protein heterodimerization activity, nucleosome, intracellular non-membrane-bounded organelle and translation were among the top enriched terms at both 3 h and 7 h. These results suggest that the Cd-responsive transcriptome was strongly associated with cellular architecture, protein-complex formation, chromatin-associated organization and translational machinery.

Underrepresented GO terms showed a different pattern. Membrane, macromolecule modification, protein kinase activity, adenyl ribonucleotide binding and intracellular membrane-bounded organelle were among the most depleted categories at both time points. These terms should not be interpreted as transcriptionally downregulated pathways, but as broad functional categories that were less represented than expected among significant Cd-responsive genes.

Overall, the direct oat GO enrichment was consistent with the individual-gene analysis, where prominent responses included cell-wall proteins, PR60 transfer-cell-associated transcripts, DUF239-containing proteins, cysteine proteinase inhibitors, nucleosome-associated proteins and translation-related genes. Because pathway enrichment using oat gene IDs was limited by sparse KEGG and Reactome annotation for A. sativa Sang, additional KEGG, Reactome and WikiPathways analyses were performed using Arabidopsis and rice homologues as supportive functional context. These homologue-based analyses showed partially overlapping pathways, while differences between Arabidopsis and rice results likely reflect annotation depth, species-specific pathway representation or both.

### Early transcriptional response at 3 h suggests targeted adjustment of transport interfaces, cellular protection and broad stress-program repression

At 3 h after Cd exposure, the low-Cd accumulating segregant showed a selective transcriptional response rather than broad activation of all stress-associated pathways. Induced transcripts included cell-wall protein precursors, PR60 transfer-cell-associated genes, DUF239-containing genes, cysteine proteinase inhibitors, nucleosome assembly protein 1-like and actin, suggesting early adjustment of cellular architecture, transport-interface functions, protein protection and intracellular organisation. In contrast, several dehydration-, pathogen-defence-, cell-wall-loosening- and ROS-associated transcripts were repressed, indicating attenuation of broader stress programmes.

This pattern was consistent with oat GO enrichment among all significant 3 h DEGs, where structural molecule activity, protein heterodimerization activity, intracellular non-membrane-bounded organelle, nucleosome and translation were among the strongest terms. Because the enrichment analysis was not direction-specific, these categories were interpreted as biological systems represented among Cd-responsive genes rather than as activated pathways.

#### Activation of cell-wall-associated and transport-interface processes

Repeated induction of cell-wall protein precursor genes was one of the clearest features of the 3 h response. Plant cell walls can contribute to heavy-metal binding and immobilisation (Parrotta et al., 2015), and cell-wall proteins participate in rapid biotic and abiotic stress responses (San Clemente et al., 2022). In developing caryopses, where cell walls also contribute to solute exchange between maternal and filial tissues, this induction may indicate early adjustment of transport-interface properties affecting Cd movement, retention or allocation, rather than direct activation of classical Cd-detoxification

#### Endosperm transfer-cell-specific PR60 genes indicate adjustment of nutrient-transfer interfaces

Several strongly induced 3 h transcripts encoded endosperm transfer-cell-specific PR60 proteins. Endosperm transfer cells are located at the maternal–filial interface and regulate nutrient movement into developing cereal grains (Thiel, 2014; Lopato et al., 2014), making PR60 induction biologically relevant in this tissue context. Although these data do not demonstrate a direct role in Cd transport, PR60 induction may indicate early adjustment of grain-filling or solute-transfer interfaces under Cd exposure, consistent with modulation of transport-interface biology rather than canonical Cd detoxification alone.

#### Cysteine proteinase inhibitors suggest early protection of cellular viability

Several strongly induced 3 h transcripts encoded cysteine proteinase inhibitors, or cystatins. These proteins regulate papain-like cysteine proteases and have been linked to plant development and abiotic-stress responses, including drought and salinity-related contexts (Li et al., 2015; Moloi and Ngara, 2023). Because heavy-metal stress can induce programmed cell death, and inhibition of papain-like cysteine proteases has been associated with increased cell viability under heavy-metal stress (Sychta et al., 2020), cystatin induction in AS131 may indicate early protection against excessive proteolysis. This supports a cautious interpretation of selective cellular protection rather than broad stress activation.

#### DUF239-containing proteins as candidate early-response genes in developing caryopsis tissue

DUF239-containing proteins, including NEP-interacting proteins, were among the most strongly induced 3 h transcripts. Although this gene family remains poorly characterised, Arabidopsis DUF239 genes have been reported to be largely associated with reproductive organs, seed development and especially endosperm expression (Vergès et al., 2023). Their repeated induction in developing oat caryopses may therefore reflect both tissue-specific developmental regulation and early Cd-responsive transcriptional adjustment. At present, these genes should be considered candidate early-response genes rather than confirmed contributors to Cd tolerance or low Cd accumulation. Nevertheless, their recurrence among the strongest induced transcripts makes them relevant candidates for future investigation in the context of grain Cd accumulation.

#### Strong fold-change genes point to chromatin, cytoskeletal and intracellular reorganization

Transcripts with the largest fold changes pointed to additional cellular processes potentially involved in the early Cd response, including nucleosome assembly protein 1-like, actin, a pathogenesis-related thaumatin superfamily protein, endoglucanase, exportin-2 and a ZZ-type zinc finger-containing protein. Together, these genes suggest possible effects on chromatin organisation, cytoskeletal dynamics, intracellular trafficking and stress-related regulation. This interpretation is consistent with reports that Cd can induce oxidative stress and DNA damage in plants, including *Vicia faba* (Lin et al., 2007), and that actin is involved in early plant responses to heavy-metal stress (Kulikova et al., 2009). The induction of a thaumatin-related transcript may also indicate recruitment of defence-associated stress-response components, as thaumatin-like proteins have been linked to plant development and abiotic and fungal stress responses (Sharma et al., 2022). The significance-ranked and fold-change-ranked transcript sets therefore appeared to capture complementary aspects of the 3 h response. The most statistically significant transcripts emphasised cell-wall functions, transfer-cell biology and grain-development-associated processes, whereas the largest fold-change transcripts highlighted chromatin maintenance, cytoskeletal reorganisation, intracellular trafficking and stress-related regulation.

#### Repression of ROS-associated and oxidative metabolism genes may limit excessive stress amplification

Several strongly repressed 3 h transcripts were associated with oxidative metabolism and ROS-linked processes, including peroxidase, polyamine oxidase and peroxisomal hydroxy-acid oxidase. Although peroxidases are often linked to abiotic stress responses, their regulation can be stress-, tissue- and genotype-dependent, and downregulation of specific peroxidase genes under osmotic stress has been reported in wheat (Csiszár et al., 2012). Polyamine oxidases can contribute to H₂O₂ production and ROS signalling during abiotic stress (Moschou et al., 2008), while polyamines have been implicated in Cd tolerance and phytochelatin-related metabolism (Pál et al., 2017). Therefore, repression of polyamine oxidase and other ROS-associated transcripts may reflect moderation of ROS-generating or ROS-amplifying pathways during the early Cd response. Rather than indicating a lack of stress response, this pattern may suggest controlled attenuation of oxidative signalling in the low-Cd segregant, potentially limiting excessive oxidative damage or unnecessary activation of programmed-cell-death-associated processes.

#### Repression of LEA, dehydrin and seed-maturation-associated genes suggests attenuation of broad dehydration programmes

Several strongly repressed 3 h transcripts encoded LEA-, dehydrin- and seed-maturation-associated proteins. LEA genes are associated with seed maturation, ABA-related regulation and abiotic-stress responses, but their expression is highly variable among gene family members, tissues and conditions (Bies-Ethève et al., 2008; Jia et al., 2022). Their coordinated repression suggests that Cd exposure did not induce a broad dehydration- or seed-maturation-type programme in AS131 at this early time point. Instead, AS131 appeared to repress general LEA/dehydrin-associated stress responses while inducing more specific interface-, protection- and caryopsis-associated transcripts, consistent with selective early regulation rather than generalized stress activation.

#### Selective reprogramming of cell-wall-related processes

Several strongly repressed 3 h transcripts were associated with cell-wall loosening or defence-related wall metabolism, including expansin-B15 and glucan endo-1,3-β-glucosidase. Expansins regulate cell-wall loosening and growth, and their expression is often tissue- and stress-dependent (Chen et al., 2019; Han et al., 2019). Their repression, together with simultaneous induction of cell-wall protein precursor genes, suggests that the 3 h response did not involve uniform activation or suppression of cell-wall processes. Instead, AS131 may have selectively adjusted cell-wall functions, favouring structural or interface-associated components while repressing wall-loosening or defence-associated wall-remodelling genes.

#### Suppression of pathogen-defence-associated pathways

Several downregulated transcripts also belonged to pathogen-defence-associated systems, including endochitinases, glucanases, thaumatin-like proteins and an avenacosidase-like β-D-glucosidase. Chitinases and β-1,3-glucanases are classical fungal-defence components, while thaumatin-like proteins belong to the PR-5 family and have been linked to both biotic and abiotic stress responses (Sharma et al., 2022). The opposite regulation of different thaumatin-related transcripts likely reflects functional diversification within this gene family rather than contradictory regulation.

The repression of an avenacosidase-like transcript is also notable because oat avenacin-related pathways contribute to preformed antifungal defence, and saponin-deficient oat mutants show compromised disease resistance (Osbourn et al., 1991; Papadopoulou et al., 1999). Thus, repression of selected pathogen-defence-associated transcripts after Cd exposure may indicate reduced investment in defence programmes that are not central to the early heavy-metal response, while other interface- and protection-associated processes are prioritised.

#### Attenuation of ABA-associated and developmental signalling components

Several additional downregulated transcripts were associated with broader stress-signalling or developmental pathways. The BURP-domain-containing protein showed similarity to RD22-like proteins, which are commonly associated with ABA, drought and salinity responses. Reduced expression of this transcript therefore supports the interpretation that the low-Cd segregant did not strongly activate a classical ABA-dependent drought-response programme following Cd exposure. Similarly, repression of vacuolar sorting receptor 1, TBP-associated factor 4B and other signalling-related transcripts may indicate temporary reduction of developmental, trafficking or regulatory processes that are not prioritized during the earliest Cd response. Although the precise functional significance of these genes remains uncertain, their coordinated repression is consistent with broader transcriptional reallocation of resources.

#### Early response model: selective interface protection rather than generalized stress activation

The 3 h transcriptome suggests a selective early response rather than broad activation of stress pathways. AS131 induced genes associated with cell-wall and transport-interface functions, transfer-cell biology, cysteine proteinase inhibition, DUF239-containing proteins, chromatin-associated organization and cytoskeletal remodelling, while repressing several dehydration-, ABA-, seed-maturation-, pathogen-defence- and ROS-associated transcripts. This pattern suggests early prioritization of cellular interface protection, cell viability and caryopsis transport-related regulation, together with attenuation of broader and potentially costly stress programmes. Rather than stronger activation of canonical Cd-detoxification pathways, the 3 h response supports a hypothesis of targeted transcriptional fine-tuning that may precede the more homeostatic acclimation pattern observed at 7 h.

### The 7 h response indicates homeostatic acclimation and attenuation of broad stress signalling

The 7 h time point was interpreted as a later transcriptional phase following the initial 3 h response. Whereas the 3 h profile emphasised cell-wall functions, transfer-cell biology, DUF239-containing proteins and cysteine proteinase inhibitors, the 7 h response appeared to shift towards homeostatic adjustment, metabolism, intracellular trafficking and attenuation of broad stress-signalling programmes.

Strongly induced 7 h transcripts included TIP aquaporins, thiamine thiazole synthases, EF-Tu proteins, coatomer-related proteins and carbohydrate-associated enzymes, while LEA/SMP/dehydrin, FRO7-like, EF-hand calcium-binding and stress-regulatory transcripts were repressed. This pattern suggests a transition from early stress perception towards a more regulated acclimation-like state.

Pathway-level analyses were consistent with this interpretation. Direct oat GO enrichment highlighted structural, nucleosome-associated and translation-related categories, while supportive Arabidopsis and rice homologue-based analyses pointed to ribosome-related functions, amino-acid and carbohydrate metabolism, hormone-associated signalling and seed/development-related pathways. These pathway results were used as functional context, not as direct evidence of pathway activation.

#### Vacuolar homeostasis and water balance

The repeated induction of tonoplast intrinsic protein 2 (TIP2) genes was a clear feature of the 7 h response, with three homologues from different oat subgenomes among the induced transcripts. TIPs are aquaporins involved in water and small neutral solute transport across vacuolar membranes (Loqué et al., 2005), and aquaporin activity has been associated with changes in membrane water permeability under Cd and other heavy-metal stresses (Przedpelska-Wasowicz and Wierzbicka, 2011). Thus, TIP2 induction may reflect adjustment of vacuolar water balance, solute distribution and intracellular homeostasis during Cd exposure, rather than direct evidence for TIP2-mediated Cd detoxification.

#### Metabolic protection and resource allocation

Several 7 h induced transcripts suggested metabolic adjustment during Cd exposure. Induction of thiamine thiazole synthase genes may reflect activation of vitamin B1-associated metabolism supporting cellular protection, while hexosyltransferase, sucrose synthase and a class I glutamine amidotransferase-like transcript point to adjustment of carbohydrate- and nitrogen-associated metabolism. This is consistent with homologue-based pathway results highlighting starch and sucrose metabolism, amino-acid metabolism and related metabolic categories.

WCI16-like/nodulin-related transcripts were also induced. Because WCI16 has been described as a cold-acclimation-induced wheat protein with LEA-like protective properties, including cryoprotective and DNA-binding activity (Sasaki et al., 2014), its induction here may indicate broader stress-acclimation or cellular-stabilisation processes rather than a Cd-specific defence pathway. Together, these patterns suggest metabolic support and resource reallocation during the 7 h acclimation-like response.

#### Maintenance of protein synthesis, chromatin integrity and cellular machinery

Several 7 h induced transcripts were associated with translation and cellular maintenance, including EF-Tu, ribosomal protein L9 and histone H4. EF-Tu/EF1A proteins have been linked to drought and salinity responses and may contribute to protein protection under stress (Shin et al., 2009; Xu et al., 2023). Their induction, together with ribosomal and histone-associated transcripts, may indicate maintenance or adjustment of protein synthesis and chromatin-related functions during Cd exposure.

This agrees with the oat GO enrichment, where translation and nucleosome-associated categories were strongly represented at 7 h, and with homologue-based pathway analyses showing ribosome-related enrichment. Thus, the 7 h response may reflect preservation of cellular machinery during acclimation rather than broad developmental shutdown.

#### Intracellular trafficking and transport regulation

Several 7 h induced transcripts were linked to intracellular transport and signalling, including coatomer beta′ subunit, auxin efflux carrier family protein, casein kinase I and response regulator ORR22. COPI coatomer complexes are involved in vesicle trafficking and intracellular protein transport, and Arabidopsis β-COP function has been associated with growth and salt-stress tolerance (Sánchez-Simarro et al., 2020). COPI-associated trafficking has also been linked to regulation of metal-uptake machinery, including IRT1 (Abuzeineh et al., 2022), a broad-spectrum transporter capable of transporting Cd (Spielmann et al., 2022). Thus, induction of coatomer-related transcripts may reflect adjustment of intracellular trafficking, transporter turnover or solute allocation during Cd exposure, rather than direct evidence for Cd transport. Induction of an auxin efflux carrier further supports a possible role for transport regulation and developmental reprogramming in the later response, although this connection remains hypothetical.

#### Repression of dehydration- and seed-maturation-associated programmes

Several 7 h downregulated transcripts encoded LEA-related proteins, including LEA EMB564, dehydrins and seed maturation proteins (SMPs). LEA proteins are associated with dehydration tolerance, seed maturation, ABA-related regulation and abiotic-stress responses, while SMPs represent a LEA subgroup linked to seed maturation and desiccation processes (Du et al., 2013; Zou et al., 2022). Their repression in developing caryopses, together with similar patterns observed at 3 h, suggests that Cd exposure did not trigger a broad LEA/dehydration-type protective programme in AS131. Instead, the response appears more consistent with attenuation of general dehydration- and maturation-associated programmes alongside induction of homeostasis-, metabolism- and trafficking-related transcripts.

#### Adjustment of redox homeostasis and metal-associated metabolism

Several strongly downregulated transcripts at 7 h were associated with redox metabolism and metal-associated homeostasis, including FRO7-like ferric reduction oxidases, polyamine oxidase and peroxisomal hydroxy-acid oxidase. Two FRO7-like transcripts were significantly repressed following Cd treatment. Because FRO proteins are linked to iron/redox homeostasis, including chloroplast-associated iron metabolism, repression of FRO7-like transcripts in developing oat caryopses may reflect adjustment of iron-related redox balance rather than direct regulation of Cd transport.

The simultaneous repression of polyamine oxidase and peroxisomal hydroxy-acid oxidase further suggests that selected ROS-linked metabolic processes were attenuated at 7 h. This interpretation is consistent with similar repression of ROS-associated transcripts at 3 h. However, because ROS levels were not directly measured, these results should be interpreted as transcript-level evidence for redox-associated adjustment rather than proof of reduced oxidative activity.

#### Fine-tuning of calcium signalling under Cd exposure

An EF-hand calcium-binding protein transcript was among the strongly downregulated genes at 7 h. EF-hand-containing proteins are important components of plant Ca²⁺ signalling networks, including calmodulins, calmodulin-like proteins, calcineurin B-like proteins and calcium-dependent protein kinases (Mohanta et al., 2019; Zeng et al., 2015). Because Cd²⁺ and Ca²⁺ can interact in plants, and Cd exposure may interfere with Ca²⁺-dependent signalling, this transcript is a relevant candidate for interpreting the later Cd response (Chen et al., 2018; Liu et al., 2023).

The transcript should nevertheless be interpreted cautiously. It likely encodes a Ca²⁺-binding signalling component rather than a Ca²⁺ channel or Cd transporter, and therefore its repression cannot be taken as evidence for altered Cd uptake or Ca²⁺ influx. Instead, it may reflect transcript-level fine-tuning of Ca²⁺-related signal decoding during the 7 h response.

#### Repression of broad stress-regulatory networks

Several 7 h downregulated transcripts encoded regulatory proteins, including a DUF1664-containing bZIP transcription factor, a MYB86-like transcription factor, casein kinase I and another protein kinase family member. Because bZIP, MYB and kinase families are broadly involved in stress signalling and transcriptional regulation, their repression may reflect attenuation or refinement of general stress-regulatory outputs rather than direct metal-detoxification activity.

The MYB86-like transcript is notable, although its function in oat is unknown. In Arabidopsis, MYB49 promotes Cd accumulation through regulation of metal-uptake genes including IRT1 (Zhang et al., 2019), but this mechanism cannot be directly inferred for oat MYB86. Its repression should therefore be interpreted cautiously as a candidate regulatory signal that may warrant further investigation in the low-Cd phenotype.

#### Reduced investment in extracellular, developmental, pathogen-defence and chloroplast-associated processes

Several 7 h downregulated transcripts were associated with extracellular, developmental or defence-related functions, including glycine-rich proteins, BURP-domain proteins, endochitinases, β-D-glucosidase and ELIP/HV90. Together, these genes suggest attenuation of selected cell-wall, dehydration/stress-associated and pathogen-defence programmes during the later Cd response.

The continued repression of endochitinase and β-D-glucosidase transcripts was consistent with the 3 h pattern, where chitinase-, glucanase- and avenacosidase-like genes were also downregulated. Because avenacosidase-like β-glucosidases contribute to activation of antimicrobial avenacin-type defence compounds in oat (Osbourn et al., 1991; Papadopoulou et al., 1999), this may indicate reduced investment in pathogen-defence pathways during early Cd acclimation. Repression of ELIP/HV90 may additionally reflect adjustment of chloroplast-associated activity, rather than a specific Cd-detoxification response.

#### Later response interpretation: attenuation of alarm signalling accompanies acclimation

Taken together, the 7 h transcriptional profile suggests a shift from the early 3 h interface/protection response towards a more homeostatic acclimation-like state. Whereas the 3 h response emphasised cell-wall-associated structures, transfer-cell biology and cellular protection, the 7 h response was more strongly associated with vacuolar homeostasis, metabolism, intracellular trafficking and protein synthesis. Induction of TIP aquaporins, thiamine biosynthesis genes, EF-Tu proteins, coatomer-related transcripts and carbohydrate-associated metabolic genes is consistent with adjustment of cellular maintenance and metabolic support during Cd exposure.

At the same time, several transcripts associated with dehydration, redox metabolism, calcium signalling, pathogen defence and broad stress regulation were repressed, including LEA/SMP/dehydrin genes, FRO7-like ferric reduction oxidases, EF-hand calcium-binding proteins, MYB/bZIP transcription factors, kinase-related transcripts, glycine-rich proteins, BURP-domain proteins and selected pathogen-defence-associated genes. This pattern suggests attenuation or refinement of broad stress-signalling outputs rather than continued escalation of a general defence response.

Importantly, these results do not indicate stronger activation of classical Cd-detoxification pathways. Instead, they support a working hypothesis in which the low-Cd segregant may regulate water balance, vacuolar function, intracellular trafficking, metabolic support and developmental signalling during early Cd exposure. This interpretation remains hypothesis-generating, but the repeated emergence of homeostasis-, transport-, metabolism- and signalling-associated processes suggests candidate biological layers for future investigation of the low-Cd phenotype.

Several top-ranked Cd-responsive transcripts also appeared to include related copies from different oat subgenomes. This observation was not tested genome-wide, but it raises the possibility that some Cd-responsive gene families may be coordinately regulated across homoeologous loci.

### Concordance between pathway-level and transcript-level responses

Pathway-level and transcript-level analyses provided complementary views of the Cd response. GO enrichment identified biological systems represented among Cd-responsive genes, whereas individual transcript analysis provided direction and biological context. At 3 h, enriched terms related to structural molecule activity, protein heterodimerization, nucleosome-associated components and translation were consistent with induction of cell-wall protein precursors, PR60 transfer-cell-associated transcripts, DUF239-containing proteins, cysteine proteinase inhibitors, nucleosome assembly protein 1-like and actin. This supports early effects on cellular architecture, transport interfaces and protein protection. In parallel, repression of LEA-related, pathogen-defence and ROS-associated transcripts suggested that broad stress programmes were not uniformly activated.

At 7 h, oat GO and supportive Arabidopsis/rice homologue-based analyses pointed to translation, ribosome-related functions, carbohydrate and amino-acid metabolism, hormone-associated signalling and seed/development-related pathways. This was consistent with induction of TIP2, thiamine thiazole synthase, EF-Tu, coatomer-related and sucrose-associated transcripts, together with repression of LEA/SMP/dehydrin, FRO7-like, EF-hand calcium-binding and stress-regulatory transcripts.

Together, these results suggest a shift from early transport-interface and cellular-protection responses at 3 h towards a more homeostatic acclimation-like response at 7 h. Because GO analyses used all significant DEGs, pathway results were interpreted as affected biological systems, while directionality was inferred from individual transcripts.

### Canonical Cd-detoxification genes were not the dominant early transcriptional signal

The strongest early Cd-responsive transcripts in AS131 were not dominated by canonical detoxification genes, such as metallothioneins, phytochelatin biosynthesis genes, heavy-metal ATPases or other classical metal-chelation components. Instead, the most recurrent signals involved transport-interface regulation, cellular protection, vacuolar and water homeostasis, intracellular trafficking, protein synthesis, metabolic support and developmental processes.

At 3 h, induced genes were mainly associated with cell-wall proteins, PR60 transfer-cell transcripts, DUF239-containing proteins and cysteine proteinase inhibitors, while several dehydration-, pathogen-defence- and ROS-associated transcripts were repressed. By 7 h, the response shifted towards TIP2 aquaporins, thiamine thiazole synthases, EF-Tu proteins, coatomer-related transcripts and carbohydrate-associated metabolism, together with repression of LEA/SMP/dehydrin, FRO7-like, EF-hand calcium-binding and stress-regulatory transcripts.

Thus, the early caryopsis response in AS131 appears more consistent with selective regulation of cellular interfaces, homeostasis, intracellular transport and developmental programmes than with broad activation of classical Cd-detoxification pathways alone. This does not exclude a role for detoxification mechanisms, but supports a hypothesis that reduced grain Cd accumulation may also depend on genotype-specific regulation of solute movement, cellular stability and caryopsis development.

### Limitations and future directions

The present results are transcriptomic and hypothesis-generating. Transcript abundance does not necessarily reflect protein abundance, enzyme activity or transporter function, and the data do not allow direct inference of Cd fluxes, tissue-specific Cd localisation or subcellular compartmentation in developing caryopses.

Because the analysis focused on one low-Cd accumulating F2 segregant, additional low- and high-Cd genotypes are needed to determine which responses are consistently associated with low Cd accumulation. Arabidopsis and rice homologue-based pathway analyses were used only as supportive functional context, not as direct evidence of oat gene function. Future work should validate candidate genes and processes related to transport interfaces, vacuolar homeostasis, intracellular trafficking, redox regulation and developmental programmes using independent expression assays, fine mapping, spatial Cd localisation and functional studies.

### Working hypothesis for the low-Cd phenotype

Together, the 3 h and 7 h transcriptomic patterns support a working hypothesis in which the low-Cd accumulating oat segregant AS131 responds to Cd exposure through selective regulation of transport-interface, homeostatic and developmental processes, rather than through broad activation of canonical Cd-detoxification pathways alone. At 3 h, the strongest responses involved cell-wall protein precursors, PR60 transfer-cell-associated transcripts, DUF239-containing proteins and cysteine proteinase inhibitors, together with repression of broad dehydration-, pathogen-defence- and ROS-associated programmes. By 7 h, the response shifted towards vacuolar homeostasis, metabolic support, protein maintenance and intracellular trafficking, with induction of TIP2 aquaporins, thiamine thiazole synthases, EF-Tu proteins and coatomer-related transcripts, while LEA/SMP/dehydrin, FRO7-like, calcium-signalling and stress-regulatory transcripts were repressed. This proposed temporal model is summarised in Fig. 4.

**Figure 4.**
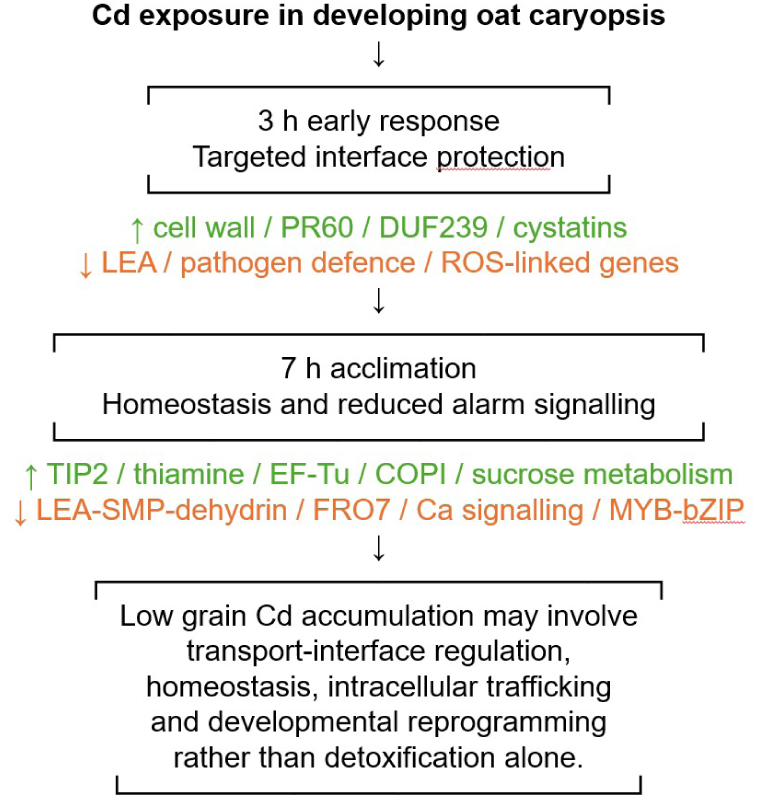
Working model of the early Cd response in the low-Cd accumulating oat segregant AS131. The model proposes that the low-Cd phenotype may involve coordinated regulation of transport interfaces, intracellular homeostasis and caryopsis developmental processes, rather than broad activation of canonical Cd-detoxification pathways alone.

This pattern suggests that reduced Cd accumulation in developing oat grain may involve genotype-specific regulation of solute movement, cellular maintenance, transport-interface activity and caryopsis developmental programmes, in addition to or partly instead of enhanced detoxification capacity alone. In durum wheat and maize, major effects on grain Cd accumulation have been linked to HMA3-type metal transporter genes, showing that individual transport or sequestration genes can strongly influence grain Cd levels (Maccaferri et al., 2019; Tang et al., 2021). However, the early oat caryopsis transcriptome described here did not show dominant activation of canonical detoxification genes. Instead, the strongest responses in AS131 pointed towards transport-interface adjustment, cellular protection, homeostatic regulation and developmental processes.

We therefore hypothesise that early transcriptional reprogramming in the developing caryopsis may help establish a cellular environment compatible with reduced Cd accumulation, potentially complementing major genetic factors controlling Cd partitioning or retention. This model remains hypothesis-generating, but it provides a testable framework for future studies of Cd allocation and low-Cd grain accumulation in oat.

## Conclusions

This study provides an early transcriptomic view of Cd responses in developing oat caryopses of a low-Cd accumulating F₂ segregant. Rather than being dominated by canonical Cd-detoxification pathways, the response was characterized by coordinated changes in transport-interface activity, cellular protection, homeostatic regulation, intracellular trafficking and developmental processes.

GO enrichment and homologue-based pathway analyses broadly supported these transcript-level observations, indicating a transition from early cellular reorganization towards a more acclimated physiological state. Collectively, the results suggest that reduced grain Cd accumulation in oat may involve regulation of solute-transfer interfaces and caryopsis homeostasis in addition to classical detoxification mechanisms.

These findings provide a hypothesis-generating framework for future studies aimed at identifying the mechanisms and genes underlying low Cd accumulation in oat grain.

## Availability of data and materials

RNA-seq data will be made available in the European Nucleotide Archive (ENA) under an accession number upon publication.

## Acknowledgements

This work was funded by the HAKA project, supported by the Ministry of Agriculture and Forestry of Finland (Diary number VN/9138/2023), and co-funded by the Natural Resources Institute Finland (Luke). We thank Marja-Riitta Arajärvi, Auli Kedonperä, Leena Holkeri and Kirsi Puisto for excellent laboratory and technical assistance, and Merja Eurola for cadmium analyses.

## Declaration of generative AI and AI-assisted technologies in the writing process

During the preparation of this work, the author(s) used Microsoft 365 Copilot to improve language and readability, and for shortening, re-structuring and rewriting. The author(s) reviewed and edited the content and take full responsibility for the content.

## Notes

### Competing Interest Statement

The authors have declared no competing interest.

